# Light sheet microscopy for everyone? Experience of building an OpenSPIM to study flatworm development

**DOI:** 10.1101/045187

**Authors:** Johannes Girstmair, Anne Zakrzewski, François Lapraz, Mette Handberg-Thorsager, Pavel Tomancak, Peter Gabriel Pitrone, Fraser Simpson, Maximilian J. Telford

**Affiliations:** Department of Genetics, Evolution and Environment, University College London, London, WC1E 6BT United Kingdom; CNRS, CBD UMR5547, Université de Toulouse, UPS, Centre de Biologie du Développement, Bâtiment 4R3, 118 Route de Narbonne, 31062 Toulouse, France; Max Planck Institute of Molecular Cell Biology and Genetics, Pfotenhauerstr. 108, 01307 Dresden, Germany

## Abstract

**BACKGROUND:** Selective plane illumination microscopy (SPIM a type of light-sheet microscopy) involves focusing a thin sheet of laser light through a specimen at right angles to the objective lens. As only the thin section of the specimen at the focal plane of the lens is illuminated, out of focus light is naturally absent and toxicity due to light (phototoxicity) is greatly reduced enabling longer term live imaging. OpenSPIM is an open access platform (Pitrone et al. 2013 and OpenSPIM.org) created to give new users step-by-step instructions on building a basic configuration of a SPIM microscope, which can in principle be adapted and upgraded to each laboratory’s own requirements and budget. Here we describe our own experience with the process of designing, building, configuring and using an OpenSPIM for our research into the early development of the polyclad flatworm *Maritigrella crozieri* – a non-model animal.

**RESULTS:** Our OpenSPIM builds on the standard design with the addition of two colour laser illumination for simultaneous detection of two probes/molecules and dual sided illumination, which provides more even signal intensity across a specimen. Our OpenSPIM provides high resolution 3d images and time lapse recordings, and we demonstrate the use of two colour lasers and the benefits of two color dual-sided imaging. We used our microscope to study the development of the embryo of the polyclad flatworm *Maritigrella crozieri*. The capabilities of our microscope are demonstrated by our ability to record the stereotypical spiral cleavage pattern of *Maritigrella* with high-speed multi-view time lapse imaging. 3D and 4D (3D + time) reconstruction of early development from these data is possible using image registration and deconvolution tools provided as part of the open source Fiji platform. We discuss our findings on the pros and cons of a self built microscope.

**CONCLUSIONS:** We conclude that home-built microscopes, such as an OpenSPIM, together with the available open source software, such as MicroManager and Fiji, make SPIM accessible to anyone interested in having continuous access to their own light-sheet microscope. However, building an OpenSPIM is not without challenges and an open access microscope is a worthwhile, if significant, investment of time and money. Multi-view 4D microscopy is more challenging than we had expected. We hope that our gained experience during this project will help future OpenSPIM users with similar ambitions.

## Introduction

Light-sheet illumination for microscopy is an old technology enjoying a dramatic recent renaissance due to introduction of selective plane illumination microscopy (SPIM) [1].The principle of SPIM is to use optics to form a thin sheet of light that passes through the specimen. Unlike a standard microscope in SPIM the objective lens is placed perpendicular to the direction of the light such that the sheet of light illuminates the specimen only at the focal plane of the lens. This has two important benefits; it eliminates scattered light from out of focus areas of the specimen providing a natural means of optical sectioning and, because only the imaged area is illuminated, the total amount of light hitting the specimen is orders of magnitude less than in conventional fluorescence microscopy meaning that photodamage/phototoxicity is enormously reduced and imaging over long periods is possible [1]. This latter benefit is of great significance for live imaging. OpenSPIM is an open access light-sheet microscopy design [2]; http://openSPIM.org; see also [3]. The OpenSPIM resource gives users step-by-step guidance for building a basic configuration of a SPIM microscope and includes appropriate open source software for image acquisition and processing such as Fiji (http://fiji.sc/Fiji), micromanager (https://www.micro-manager.org/), multiview reconstruction plugins [4, 5] deconvolution [6] and big data viewer (http://fiji.sc/BigDataViewer). The design can be adapted and upgraded according to the users specific requirements and budget. We have designed an OpenSPIM microscope capable of dual-sided illumination (the so called T-configuration proposed on the OpenSPIM wiki). The microscope was built following instructions from the website http://openSPIM.org with modifications required to extend the capabilities of the basic single sided illumination described there (Figure 1).

**Figure 1.**
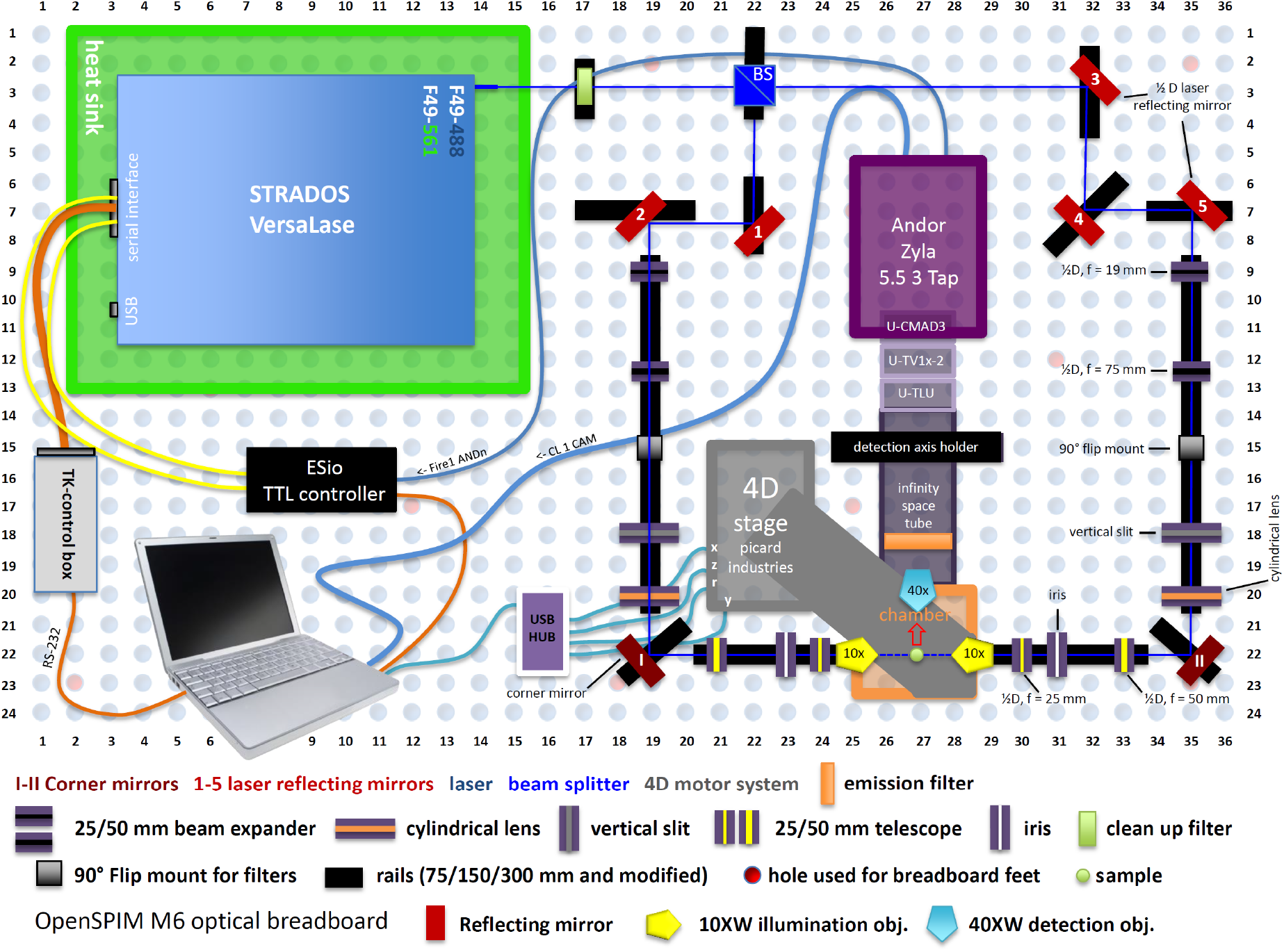
OpenSPIM with dual-sided illumination, hardware-controlled laser triggering and all hardware components

To test our system we have imaged the early embryogenesis and the larval stage of the polyclad flatworm *Maritigrella crozieri*, a promising new evo-devo model within Plathyhelminthes [7] and to study spiralian development. This polyclad flatworm has a ciliated planktotrophic larval stage known as Müller larva that shows morphological similarities to spiralianotrophic larval stage found in marine annelids and molluscs and undergoes the stereotypical spiral cleavage. The embryonic and post-embryonic development of *M. crozieri* has been previously described by [8]. Recent flatworm phylogenies confirm the basal position of polyclad flatworms within the rhabditophoran Platyhelminthes [9, 10] making *M. crozieri* and other polyclad flatworms an interesting system for evo-devo studies within Platyhelminthes and amongst other Lophotrochozoa. Our results here on both live and fixed material show that we were able to visualize the stereotypical spiral cleavage pattern of *Maritigrella crozieri* with high-speed time-lapse sequences and were able to 3D-reconstruct a number of individual time points of the early embryonic development using Fiji’s bead based registration software and multi-view deconvolution plugins [4, 6].

In this report we describe a real life experience of building an OpenSPIM microscope. We discuss the difficulties we encountered, the real costs involved including the time and effort as well as describing the limitations and significant benefits of the system.

## METHODS

### OpenSPIM - summary

An OpenSPIM microscope capable of dual-sided illumination (T-configuration) was built on an 600 × 900 × 12.7 mm aluminium breadboard following instructions from the website http://openspim.org/. The principal components of our microscope comprise a multiple wavelength laser system (Stradus VersaLase™ from Laser2000 http://www.laser2000.co.uk/versalase.php) producing two individual wavelengths (λ = 488 and 561 nm); a Zyla 5.5 3 Tap sCMOS camera from Andor (http://www.andor.com/); and a USB 4D-stage from Picard Industries (http://www.picard-industries.com/). The acquisition chamber (designed by PGP and manufactured by Pieter Fourie Design and Engineering CC; http://www.pfde.co.uk) includes openings for two 10x illumination objectives (Olympus UMPLFLN10xW left and right) and one aperture for a 40x acquisition objective (Olympus; LUMPLFLN40xW). The optical breadboard, rails and rail carriers, optical elements and mirrors were purchased from Thorlabs (http://www.thorlabs.com/), fluorescence clean up and emission filters from AHF (http://www.ahf.de/). The appendix includes a complete list of all purchased parts (Table 1) and a summary of the costs (Table 2).

**Table 1.**

| **List of quantity and materials used for building the OpenSPIM** |
| --- |
| **Laser2000**<br>Stradus VersaLase™ VersaLase 488/561<br>Heat sink (special modification) |
| **Pieter Fourie Design and Engineering CC**<br>2x RC1 vertical slit stilt<br>11x RC1 Ø1/2" lens stilt<br>3x Metal objective holder ring<br>1x Detection axis holder, base<br>1x Detection axis holder, top<br>1x Infinity space tube<br>2x Ø1"/Ø25.4 mm microscopy fluorescence emission filter holder, base<br>2x Ø1"/Ø25.4 mm microscopy fluorescence emission filter holder, top<br>8x RAIL CARRIER 15.4 mm, MOD ONLY<br>1x Acrylic sample chamber T, OLYMPUS<br>1x Metal chamber holder T, OLYMPUS<br>8x INSERT FOR RAIL CARRIER 15.4 mm (RC1 MODIFIED)<br>2x RC1 MOD, Ø1/2" lens stilt<br>2x RC1 Iris stilt<br>5x RC1 Ø1/2" mirror stilt |
| **AHF Fluorescent filters**<br>1x F72-866; 446/523/600/677 HC Quadband Filter (Emission Filter)<br>1x F59-486; Dual Line Laser Clean-up ZET 488/561 |
| **Picard Industries**<br>1x USB-4D-STAGE |
| **Thorlabs parts**<br>2x DG05-1500-H1-MD; Ø1/2" SM05-Mounted Frosted Glass Alignment Disk w/Ø1 mm Hole<br>2x NE20A-A; Ø25 mm AR-Coated Absorptive Neutral Density Filter, SM1-Threaded Mount, 350-700 nm, OD: 2.0<br>5x TRF90/M; 90° Flip Mount for Ø1" Filters and Optics, Metric<br>2x VA100/M; Adjustable Mechanical Slit, Metric<br>10x LMR05/M; Lens Mount for Ø1/2" Optics, One Retaining Ring Included, M4 Tap<br>5x KM05/M; Kinematic Mount for Ø12.7 mm Optics, Metric<br>2x GM100/M; Ø25.4 mm Gimbal Mirror Mount, Metric, One Retaining Ring Included<br>2x RSP1X15/M; Metric Rotation Mount, 360° Continuous or 15° Indexed Rotation<br>2x BB1-E02; Ø1" Broadband Dielectric Mirror, 400-750 nm<br>5x BB05-E02; Ø1/2" Broadband Dielectric Mirror, 400-750 nm<br>2x AC127-050-A-ML; f=50 mm, Ø1/2" Achromatic Doublet, SM05-Threaded Mount, ARC: 400-700 nm<br>2x AC127-025-A-ML; f=25 mm, Ø1/2" Achromatic Doublet, SM05-Threaded Mount, ARC: 400-700 nm<br>2x AC127-019-A-ML; f=19 mm, Ø1/2" Achromatic Doublet, SM05-Threaded Mount, ARC: 400-700 nm<br>2x AC127-075-A-ML; f=75 mm, Ø1/2" Achromatic Doublet, SM05-Threaded Mount, ARC: 400-700 nm<br>2x ACY254-050-A; f = 50 mm, Ø1" Cylindrical Achromat, AR Coating: 350 - 700 nm<br>24x RC1; Rail Carrier, 1" x 1", 1/4" (M6) Counterbored Mounting Hole<br>2x LMR1/M; Lens Mount for Ø1" Optics, One Retaining Ring Included, M4 Tap<br>2x SM1D12D; Ring-Activated SM1 Iris Diaphragm |
| 1x MB6090/M; Aluminum Breadboard, 600 mm x 900 mm x 12.7 mm, M6 Taps<br>3x AV2/M; Sorbothane Feet, M6 Thread, 20 - 32 kg (44 - 70.4 lb) Load, 4 Pieces<br>3x RLA300/M; Dovetail Optical Rail, 300 mm, Metric<br>5x RLA150/M; Dovetail Optical Rail, 150 mm, Metric<br>2x HW-KIT1/M; M4 Cap Screw and Hardware Kit<br>1x HW-KIT2/M; M6 Cap Screw and Hardware Kit<br>1x SPW602; SM1 Spanner Wrench, Graduated, Length = 3.88"<br>1x BS004; 50:50 Non-Polarizing Beamsplitter Cube, 400 - 700 nm, 1/2"<br>1x BS127CAM; 12.7 mm (0.50") Beamsplitter Cube Adapter for Compact 30 mm Cage Cube<br>1x CM1-4ER/M; Compact Clamping 4-Port Prism/Mirror 30 mm Cage Cube, M4 Tap<br>3x CL3/M; Compact Variable Height Clamp, M6 Tapped<br>1x PH30/M; Post Holder with Spring-Loaded Hex-Locking Thumbscrew, L = 30 mm<br>1x TR40/M; Ø12.7 mm x 40 mm Stainless Steel Optical Post, M4 Stud, M6-Tapped Hole |
| **Olympus**<br>2x N2667500; UMPLFLN10XW objective<br>1x N2667700; LUMPLFLN40XW objective |
| **Video camera mounts & adapters**<br>1x U-TLU single port tube with lens<br>1x U-TV1x video camera adapter (projection lens)<br>1x U-CMAD3 video camera mount adapter |
| **Andor**<br>1x Camera Zyla 5.5 3 Tap ex-demo model |
| **www.esimaging solutions.com**<br>1x ESio TTL Controller |
| **Misco.co.uk**<br>1x LN47340; Drobo 5D 5 Bays DAS Thunderbolt x2 (10Gbs x2)<br>5x LN46168; Red WD30EFRX 3TB HDD |

**Table 2.**

| **openSPIM Telford lab expenses (rounded; inc. VAT)** |  |
| --- | --- |
| Laser 488/561 | £18,520.00 |
| Self-made parts | £2,000.00 |
| Fluorescence filters | £880.00 |
| USB 4D-stage | £3,240.00 |
| Breadboard and optical elements | £5,400.00 |
| Objectives and camera mount | £4,500.00 |
| Camera | £6,500.00 |
| TTL control box | £270.00 |
| Processing computer | £4,400.00 |
| Data storage (15TB) | £1,100.00 |
| **total** | **£46,810.00** |

### OpenSPIM - assembly

Mirror components, optical elements and the acquisition chamber of the OpenSPIM were assembled and mounted on rail carriers as described in the video guide on the OpenSPIM website and is summarized below in 14 simplified steps and schematically represented in supplementary Figure 1.

Assembly steps: **Step 1:** Installation of breadboard feet onto the optical breadboard and placing it in its final position; **Step 2:** Installation of laser heatsink on the optical breadboard and fixing the laser system (VersaLase) on top of the heatsink; **Step 3:** Cutting of optical rails for corner mirrors and two reflecting mirrors and installation of the rail system onto the optical breadboard; **Step 4:** Installation of the pre-assembled acquisition chamber onto the corresponding rail; **Step 5:** Installation of the beam splitter (BS004, Thorlabs), which was placed into a cube adaptor (BS127CAM, Thorlabs), then fixed in place by a cage holder (CM1-4ER/M, Thorlabs) and finally mounted on a rail carrier; **Step 6:** Installation of all corner and laser reflecting mirrors; **Step 7:** Installation of the detection axis holder, infinity space tube, camera and its corresponding connection adapter units to the infinity space tube (U-CMAD3, U-TV1x-2 and U-TLU); **Step 8:** Installation of optical elements (beam expanders, telescope); **Step 9:** Installation of clean-up and emission filters; **Step 10:** Installation of Picard 4D stage in its correct position; **Step 11:** Connecting the controller boxes (Esio TTL controller box & VersaLase control box), VersaLase, Camera, USB 4D-stage and connecting them to the acquisition computer. The Versalase laser system was connected with the help of a controller box and a conventional RS-232 cable to the acquisition computer. ESio’s TTL controller box was connected directly to the VersaLase (not to the controller box) via individual SMB connector cables; **Step 12:** Setting up acquisition computer (installation of MicroManager and necessary plugins as well as drivers for VersaLase, Andor camera, Esio TTL controller box and pixel size calibration). The 5 second default security delay of the VersaLase laser system was disabled in the terminal of the Stradus VersaLase GUI software as described on the MicoManager website (https://micro-manager.org/wiki/Versalase); **Step 13:** Hardware configuration with Micromanager and testing for hardware recognition; **Step 14**: Final alignment of light-sheets by illuminating agarose containing fluorescent beads and/or stained specimens.

### OpenSPIM - alignment of illumination paths along the rails

The two illumination paths were aligned along the optical rails using alignment disks (DG05-1500-H1-MD, Thorlabs) and ring-activated iris apertures (SM1D12D, Thorlabs). Fine-tuning of the light paths was applied by adjusting the Kinematic Mounts (KM05/M, Thorlabs) of the laser reflecting mirrors. Note that it is important to think of appropriate laser safety measures during laser adjustments. For example we use special laser safety eyewear (from laservision) and avoid wearing reflective objects.

### OpenSPIM - alignment of the light-sheet for imaging

The excitation light-sheets generated from each illumination path (left and right) were initially aligned using the 25 mm and 50 mm telescope lenses and the adjuster knobs of the two gimbal mounts of each corner mirror (Horizontal & Vertical). Therefore the light-sheets were visualized in a column of agarose within the water filled acquisition chamber and using the lowest laser power (1mW) (Figure 2, a and b). Additionally during the alignment emission filters and cylindrical lenses were removed. By adjusting the distance between the 25 and 50 mm telescope lenses and their distance to the illumination objective, the light-sheet first appears as an indistinct broad fuzzy beam crossing the field of view horizontally from left to right (Figure *2*, a, orange arrows). The telescope lenses can then be further adjusted to increase the sharpness of the beam (step 1 in Figure *2*, a) and to centre its focal point (step 2 in Figure *2*, a). Next the horizontal gimbal mount adjuster knob is adjusted to bring the light-sheet in focus with the detection objective up to the point where it can be seen as a very thin stripe instead of a coarse beam. Finally the vertical gimbal mount adjuster knob is adjusted to center the light-sheet (step 3 in Figure *2*, b). In this way both illumination paths are aligned and carefully centered until they overlap each other. By putting the cylindrical lenses and the emission filter back, the alignment of both excitation light-sheets can now be tested on fluorescent beads, which ideally homogenously cover the field of view and/or on a specimen embedded in agarose. Usually additional fine tuning by adjusting the horizontal gimbal mount knob once more is necessary to achieve best imaging results.

**Figure 2.**
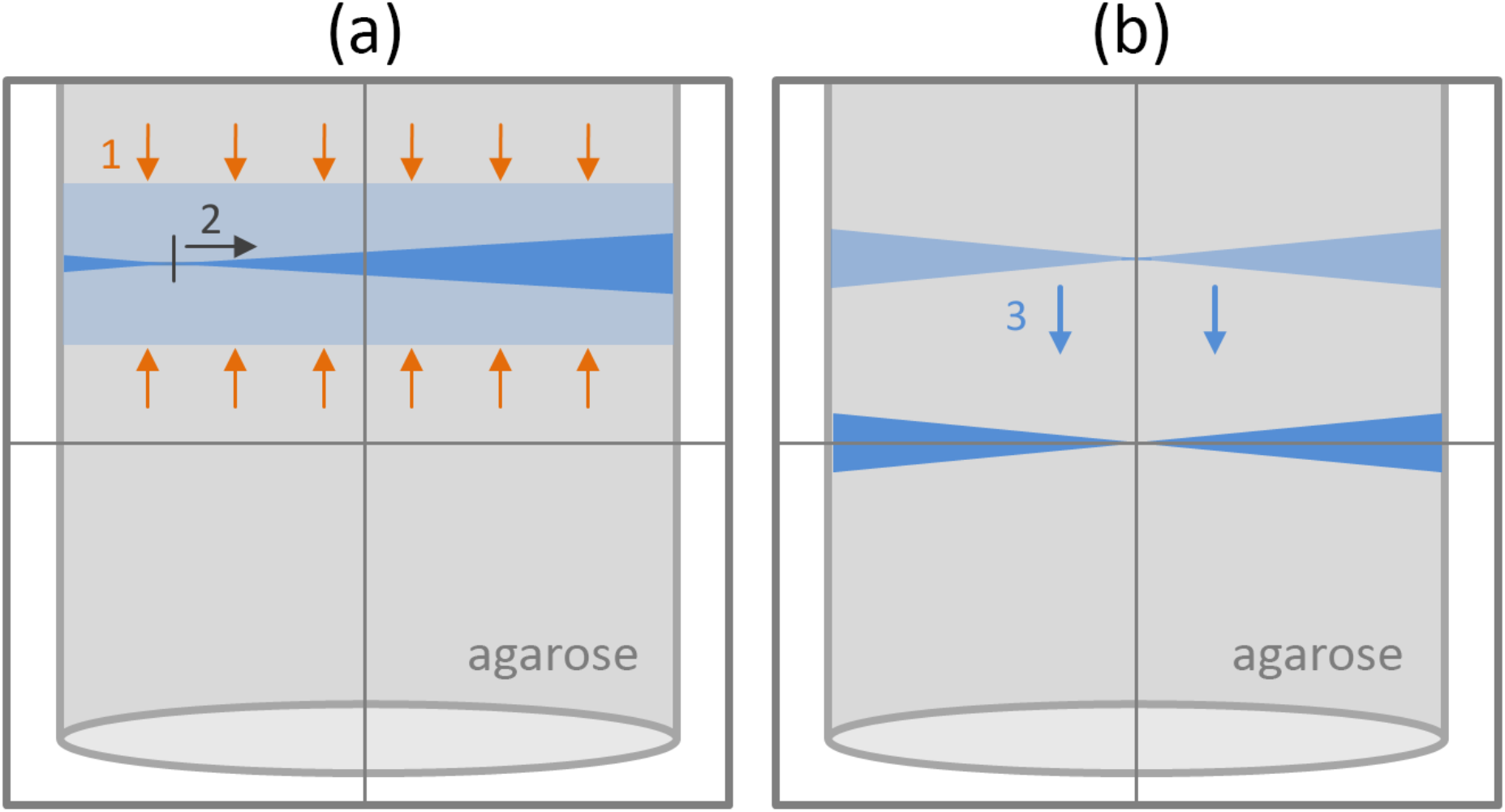
(a) and (b) Schematic drawing of laser beam visualized on agarose hanging from above into the water filled acquisition chamber. Also seen are three alignment steps of the laser beam (1-3). (a) The initially visible fuzzy beam is indicated by a bright blue horizontal stripe in between orange arrows. This coarse beam is then brought into focus with the detection objective (step1) and therefore appears as a much thinner laser beam indicated by a blue horizontal stripe in an hourglass-like shape. Note that in this example the focal point of the beam is at this point still shifted to the left (vertical grey line) and need further adjustments (step2). (b) The laser beam is shifted from the top to a central position within the field of view (step3).

### OpenSPIM - configuration of the acquisition computer

All necessary hardware component drivers were installed on a HPZ820 workstation computer (see Table 3 in the appendix for computer specifications) and the OpenSPIM hardware configured with the open source microscopy software MicroManager (version 1.4.19; November 7, 2014 release; https://www.micro-manager.org/).

**Table 3.**

|  |
| --- |
| **Acquisition and processing computer information** |
| **Product:** HP Z820 Workstation |
| **Processor:** 2x Xeon E5-2630 v2 2.60Ghz |
| **Drives:** 1x 256GB SSD; 3x 3TB Hard drives |
| **Graphics:** 2x Nvidia Quadro K4000 graphic cards |
| **Memory:** 128GB RAM |

### OpenSPIM - processing of acquired data

Post-processing of acquired data was performed with the latest version of the freely available imaging software Fiji [11]. For the 3D reconstructions, we took advantage of the bead based registration algorithm and the multi-view deconvolution plugin [4-6].

### Animal culture

Adult specimens of *M. crozieri* were collected in coastal mangrove areas in the Lower Florida Keys, USA in November 2014. Eggs without egg-shells (to produce ‘naked’ embryos) were obtained from adults by poking with a needle (BD Microlance 3) and raised in Petri dishes coated with 2% agarose (diluted in filtered artificial seawater) or gelatin coated Petri dishes at room temperature in penicillin-streptomycin (100 μg/ml penicillin; 200 μg/ml streptomycin) treated Millipore filtered artificial seawater (35-36 ‰).

### In vitro synthesis of mRNA

The plasmids carrying the nuclear marker pCS2-H2B-GFP (GFP-Histone) and the surface marker pDestTol2pA2-CAAX-EGFP [12] were linearized with the restriction enzymes NotI and BglII respectively. Ambion’s SP6 mMESSAGE mMACHINE kit was used to produce capped mRNA.

### Microinjections

Fine-tipped microinjection needles were pulled on a Sutter P-97 micropipette puller (parameters: P=300; H=560; Pu=140; V=80; T=200.) and microinjections of synthesized mRNA (~300-400 ng/μl per mRNA in nuclease-free water) were carried out under a Leica DMI3000 B inverted scope with a Leica micromanipulator and a Picospitzer^®^ III at room temperature.

### Time-lapse image acquisition & imaging of embryos for 3D-reconstructions

Live embryos were briefly incubated in 0.2% low melting agarose and immediately sucked into fluorinated ethylene propylene (FEP) tubes (Bola S1815-04), which were mounted into the OpenSPIM chamber via a 1 ml BD Plastikpak (REF 300013) syringe. The use of FEP tubes has been previously described [13] and allows for the use of lower percentage agarose, thus perturbing embryo growth and development less. To capture high-speed time-lapse videos of early quartet formation at the start of development (3-4 cell stage), time-points were captured every 90 seconds. The interval between images at later stages was gradually increased from 2-3 min (4-8-cell stage), 4 min (8-128-cell stage) and finally every 7 min. For 3D reconstructions of fixed embryos, glass capillaries were used instead of FEP tubes and embryos were embedded in 1% agarose containing 0.5 μm sized fluosphere beads (1:2500, F8813 from Life Technologies) to enable registration of images taken from different angles. In this case, the agarose is pushed out of the glass capillary once mounted onto the OpenSPIM to a point that the embryo is visible outside the capillary for optimal imaging.

### Fixation and imaging of embryos used for scanning electron microscopy (SEM)

Batches of embryos were raised until development reached the desired stage (1-cell, 2-cell, 4-cell, 8-cell, 16-cell, 32-cell, 64-cell, 128-cell and intermediate phases). Fixation was done at 4°C for 1 hour in 2.5% glutaraldehyde, buffered with phosphate buffered saline (PBS; 0.05 M PB/0.3 M NaCl, pH 7.2) and post-fixed at 4°C for 20 min in 1% osmium tetroxide buffered with PBS. Fixed specimens were dehydrated in an ethanol series, dried via critical point drying, and subsequently sputtered coated with carbon or gold/palladium in a Gatan 681 High Resolution Ion Beam Coater and examined with a Jeol 7401 high resolution Field Emission Scanning Electron Microscope (SEM).

### Immunohistochemistry

1 day old larvae were relaxed for 10 to 15 min in 7.14% MgCl2 * 6H_2_O and fixed for 60 min in 4% formaldehyde (from 16 *%* paraformaldehyde: 43368 EM Grade, AlfaAesar) in 0.1 M phosphate buffer saline (PBS) at room temperature or at 4°C overnight, followed by a 5 times washing step in PBS. The larvae were subsequently stepwise transferred into 100% methanol (25%, 50%, 75%, 2x 100%) and stored at -20°C. Embryos were fixed in the same way but without the MgCl_2_ relaxation step.

Larvae and embryos were rehydrated from methanol to 0.1% Triton X-100 in 0.1 M phosphate-buffered saline (PBST) by four PBST washing steps, each reducing the concentration of methanol in PBST by 25%. Larvae (not embryos) were subsequently treated with proteinase K (0.1 mg/ml in PBST) for 8 minutes and quickly rinsed several times in PBST. Two drops of Image-iT™FX Signal Enhancer (Molecular Probes) were added to specimens, followed by four PBST washes (5 min each) and a 2-hour blocking step in 1% bovine serum albumin diluted in PBST (BSA solution). Primary antibody (1:250 monoclonal Mouse anti-Acetylated Tubulin antibody from Sigma, which labels stabilized microtubules and ciliated cells) and a secondary antibody (1:500 Alexa Fluor^®^ 568 Goat anti-Mouse from Invitrogen™) were diluted in BSA solution. Primary antibody incubation took place at 4°C overnight in the dark, followed by several washes of PBST. Then secondary antibody incubation took place at 4°C overnight in the dark, followed by several washes of PBST. Additionally 0.1 uM of the nuclear stain SytoxGreen (Invitrogen) was added during the final wash to specimens for 30 min and rinsed with PBST for 1 hour.

## RESULTS

### Our OpenSPIM produces high quality images which we compare to scanning electron micrographs (SEM)

To test whether our OpenSPIM microscope can produce high quality images, we stained fixed 1 day old Müller’s larvae with monoclonal Mouse anti-Acetylated Tubulin antibody (Sigma) and used a secondary antibody conjugated to Alexa Fluor^®^ 568 Goat anti-Mouse (Invitrogen™). Our OpenSPIM images (shown as maximum projections) show cilia covering the whole epidermis of the polyclad larva (Figure 3, A-E). This dense film of short cilia can easily be distinguished from longer cilia comprising the ciliary band along the eight lobes (Figure 3, E). Our OpenSPIM images show a clear resemblance to scanning electron microscopy images [7] of similar stage larvae (Figure 3 D, E and F) confirming reliable image acquisition with OpenSPIM at the level of embryo morphology. For a more detailed comparison of several views see also supplementary Figure 2 (A-C and A’-C’).

**Figure 3.**
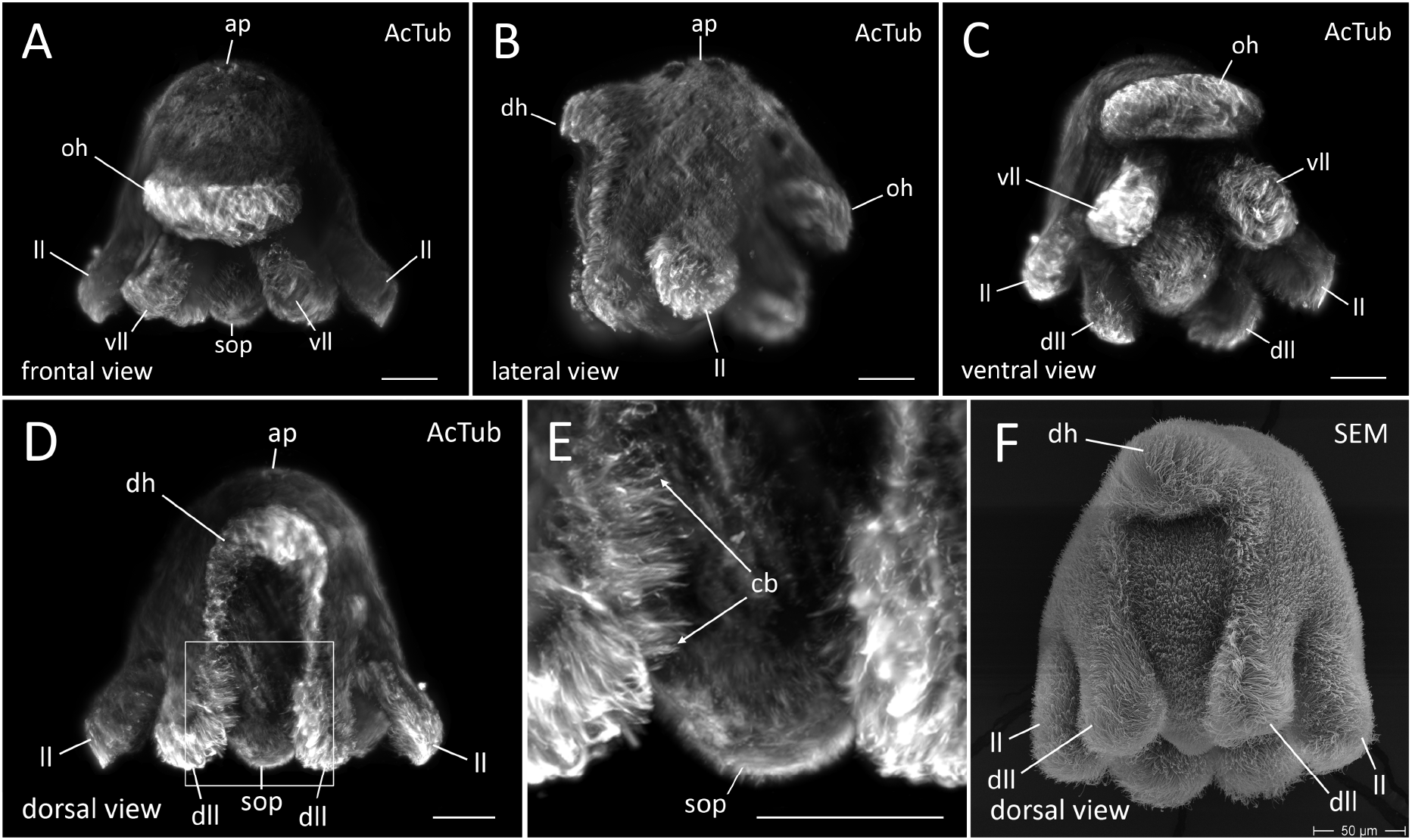
Maximum projections of fixed Müller’s larvae stained with Acetylated tubulin and imaged with OpenSPIM (A-E). (A) Anterior view (B) Lateral view (C) Ventral view (D) Posterior view (E) magnified view of area boxed in D (F) Posterior view of a *M. crozieri* larval stage obtained with scanning electron microscopy for comparison with imaging acquired by our OpenSPIM. Ap, apical plate; oh, oral hood; vll, ventro-lateral lobe; ll, lateral lobe; dll, dorso-lateral lobe; sop, sub-oral plate; cb, ciliary bands (long cilia). All scalebars are 50 μm.

### The advantage of OpenSPIM multi-view reconstructions over confocal microscopy and single image in *M. crozieri*

Standard confocal microscopes lack the possibility of multi-view imaging and reconstruction. This matters for the study of *M. crozieri* larvae and embryos due to the light intensity attenuation meaning we can only visualize one side of the embryo. We have found that the opacity causes significant signal loss, which becomes especially obvious in *Maritigrella* when the confocal z-stack of an imaged specimen is rotated. We demonstrate this here in a fixed larva labeled with the nucleic-stain SytoxGreen (Figure 4, right).

The second drawback of confocal is the necessity of using a slide with coverslip, which tends to cause deformation of our topologically complex larvae. The rotation of a confocal imaged larva reveals the slightly squeezed body shape of the larva. Confocal z-stacks are not suitable for further image processing (e.g. image pattern registration as described by [14, 15]

In contrast to confocal imaging, OpenSPIM offers a multi-view reconstruction method, whereby different angles of the same specimen can be fused into a single z-stack as shown in Fig. 4 (left side). The reconstructed larva not only keeps its natural shape, but also includes the information obtained from each individual angle, resulting in a whole-mount containing high-resolution signal from all sides. For *Maritigrella* larvae and embryos multi-view reconstructions achieved with OpenSPIM create a crucial advantage over confocal microscopy.

We further visualized to what extend average fusion and deconvolution [6] can improve results over a single view in an *M. crozieri* embryo when imaged with our OpenSPIM. This is relevant e.g. for early cleavage observations, when nuclei of the macromeres shift from the animal pole towards the vegetal pole of the embryo and are thus difficult to see.

In terms of imaging the entire embryo, it is not surprising that we found a clear benefit gained by applying multi-view imaging (average fusion or multi-view deconvolution of 5 angles) over a single one angle view. The differences are shown in Figure 5 (A-C); in the simple one angle view the small macromeres (cells A-D) at the vegetal extreme of the embryo are not visible (Figure 5, A).

**Figure 4.**
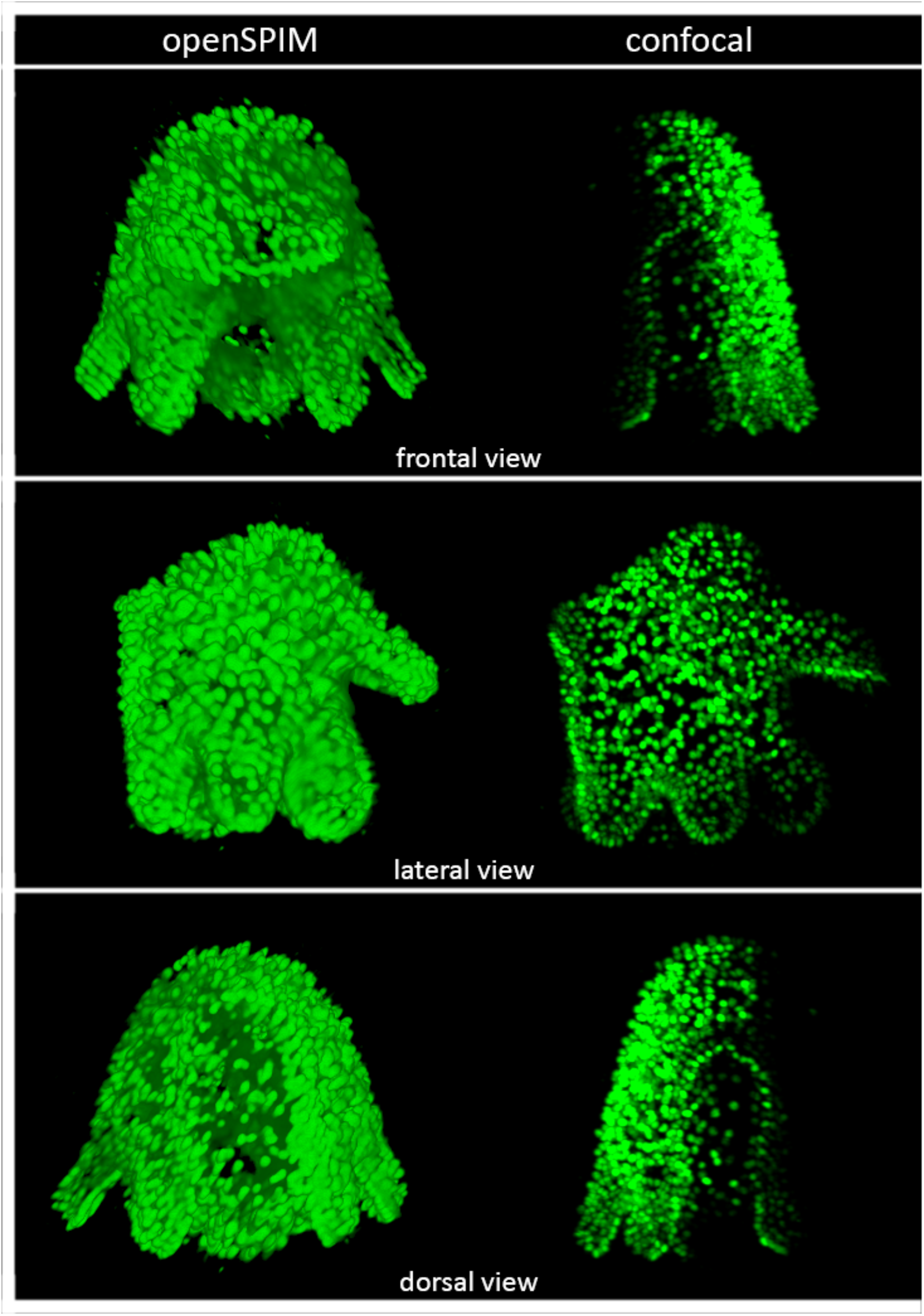
A comparison of a multi-view reconstructed larva (multi-view deconvolution of several angles) stained with the nucleic marker SytoxGreen (left side) with a larva with the same staining captured with a Leica TCS SP8 confocal laser microscopy (right side).

The acquisition of several angles (Figure 5, B and C) reveals the missing cells and makes clear that embryonic 3D reconstructions, which should include all nuclei information, depend on multi-view imaging. A slight improvement of multi-view deconvolution with 12 iterations over average fusion could be achieved (Figure 5, B and C), but appears to be less critical for early staged *M. crozieri* embryos.

**Figure 5.**
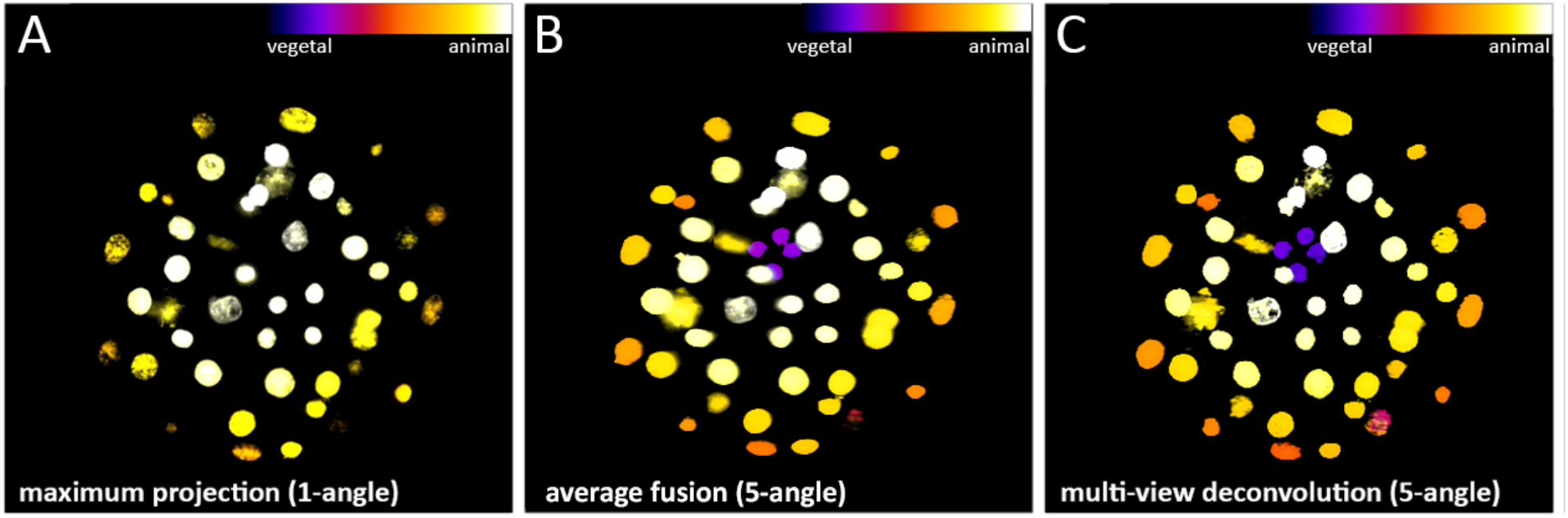
The benefit gained by applying the multi-view deconvolution method (5-angles used) vs. simple maximum projections of an acquired stack (1-angle) on an early staged *M. crozieri* embryo. **(A)** Depth color coding (Fire) of all nuclei after maximum projection of a raw z-stack (1-angle) **(B)** Depth color coding (Fire) of all nuclei after a maximum projection of a raw z-stack (1-angle) **(C)** Maximum projection of all nuclei after multi-view deconvolution. Additionally all nuclei missing in the single angle maximum projection (from B) have been coloured in red.

### Dual-sided illumination efficiently compensates axial intensity attenuation in semi-transparent specimens

Dual-sided illumination for OpenSPIM microscopy can be achieved by building a so-called T-configuration, whereby the laser beam gets split into two beams and a second optical path is installed on the optical breadboard. This allows the illumination of specimens from two sides, instead of one, as demonstrated originally by [16], and was also suggested as a potential extension in the original OpenSPIM publication [2]. The benefit of having dual-sided illumination in our OpenSPIM was tested on fixed *M. crozieri* embryos stained with the nucleic acid marker SytoxGreen. In single-sided illumination images (where one of the two illumination paths has been completely obscured - left or right respectively), a significant loss of signal during acquisition on the side of the missing illumination path becomes obvious due to axial intensity attenuation caused by our opaque and yolky specimens (Figure 6, A and C). The light attenuation is especially apparent when single nuclei from opposed illumination sites (left and right) are directly compared to each other (Figure 6, A and C, insets). In contrast, a more complete picture of the stained nuclei is achieved by using both illumination paths simultaneously (Figure 6, C, insets). This simple test clearly demonstrates the benefit of using dual-sided illumination for our slightly opaque endolecithal polyclad embryos.

**Figure 6.**
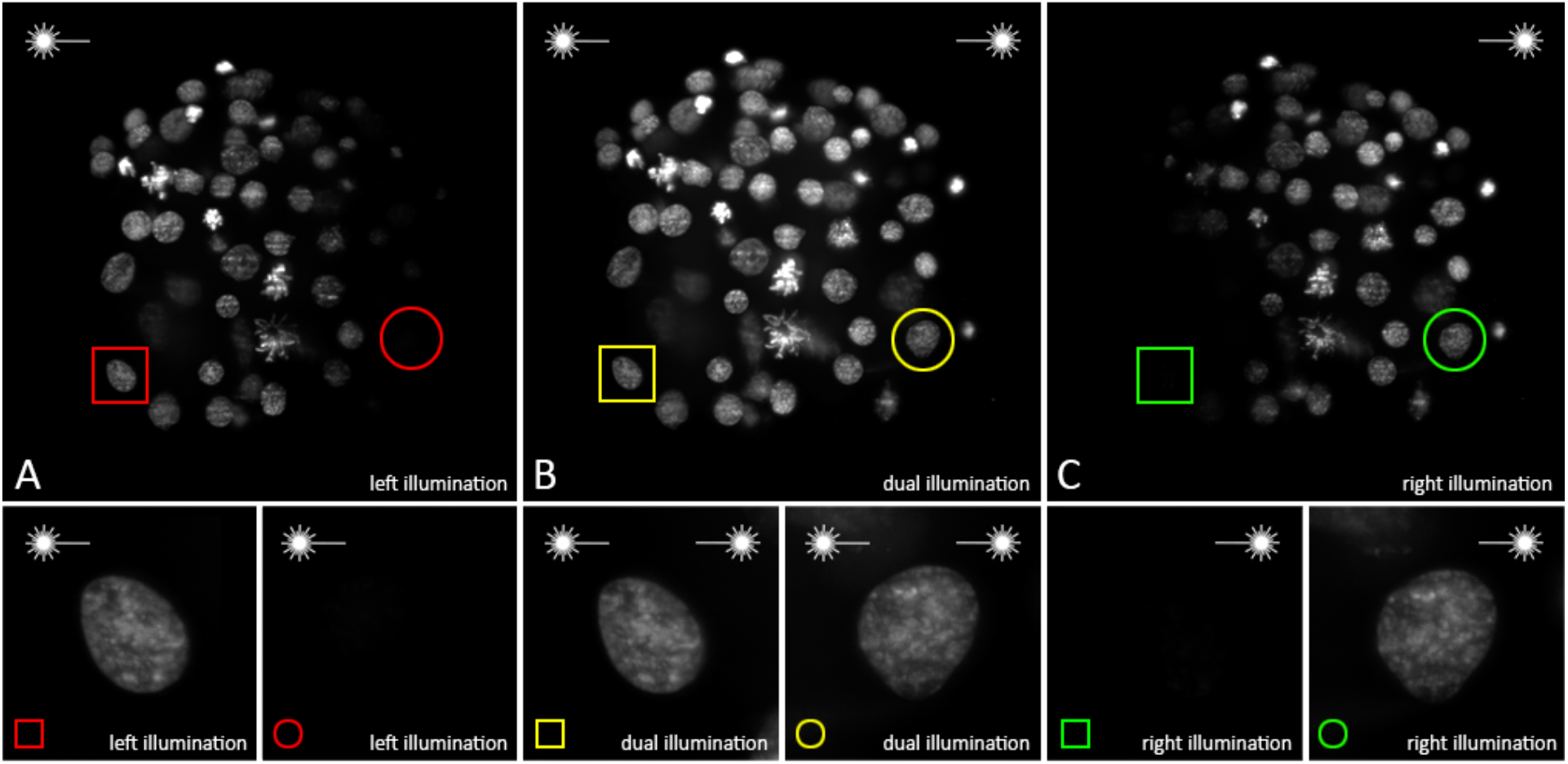
A single and dual-sided illumination, imaging test showing maximum projections of nuclei stained with the nucleic acid marker SytoxGreen. **(A)** Maximum projections of embryonic nuclei using left illumination path of the OpenSPIM microscope, **(B)** Both illumination paths and **(C)** Right illumination paths respectively. Image stacks were acquired in the following order: left illumination, right illumination, dual-sided illumination (A following C following B). All stacks and related insets have been processed identically.

### Two laser lines allow the visualization of two detection channels

Our OpenSPIM is equipped with two individual lasers (λ = 488 nm and 561 nm) that allow visualization of two detection channels. The twin laser system was tested on fixed 1 day old *M. crozieri* larvae stained with the nucleic marker SytoxGreen (488). The 561 laser was used in the same specimens to visualize auto-fluorescence of gland cells (rhabdites). The larvae have gland cell scattered mostly around the apical plate and on the ventral side of the animals, which is shown in Figure 7 in a single specimen, in which both channels (green and red) have been combined. Here we simply demonstrate the use and precise alignment of both laser beams (488 nm and 561 nm). The latter is a requirement during multi color imaging to obtain good quality images from both channels.

The initial laser beam alignments in a multiple laser system (in our case VersaLase) is done by the manufacturer and an OpenSPIM user can later only align one wavelength, e.g. 488 nm, while the other wavelength(s) (in our case the 561 nm) is presumed to coincide. It is worth noting that we have transported our OpenSPIM, allowing us to participate in courses (e.g. during the MAMED EMBO practical course in July 2015) and that the default alignment of our laser system alignment has proven robust during travelling.

**Figure 7.**
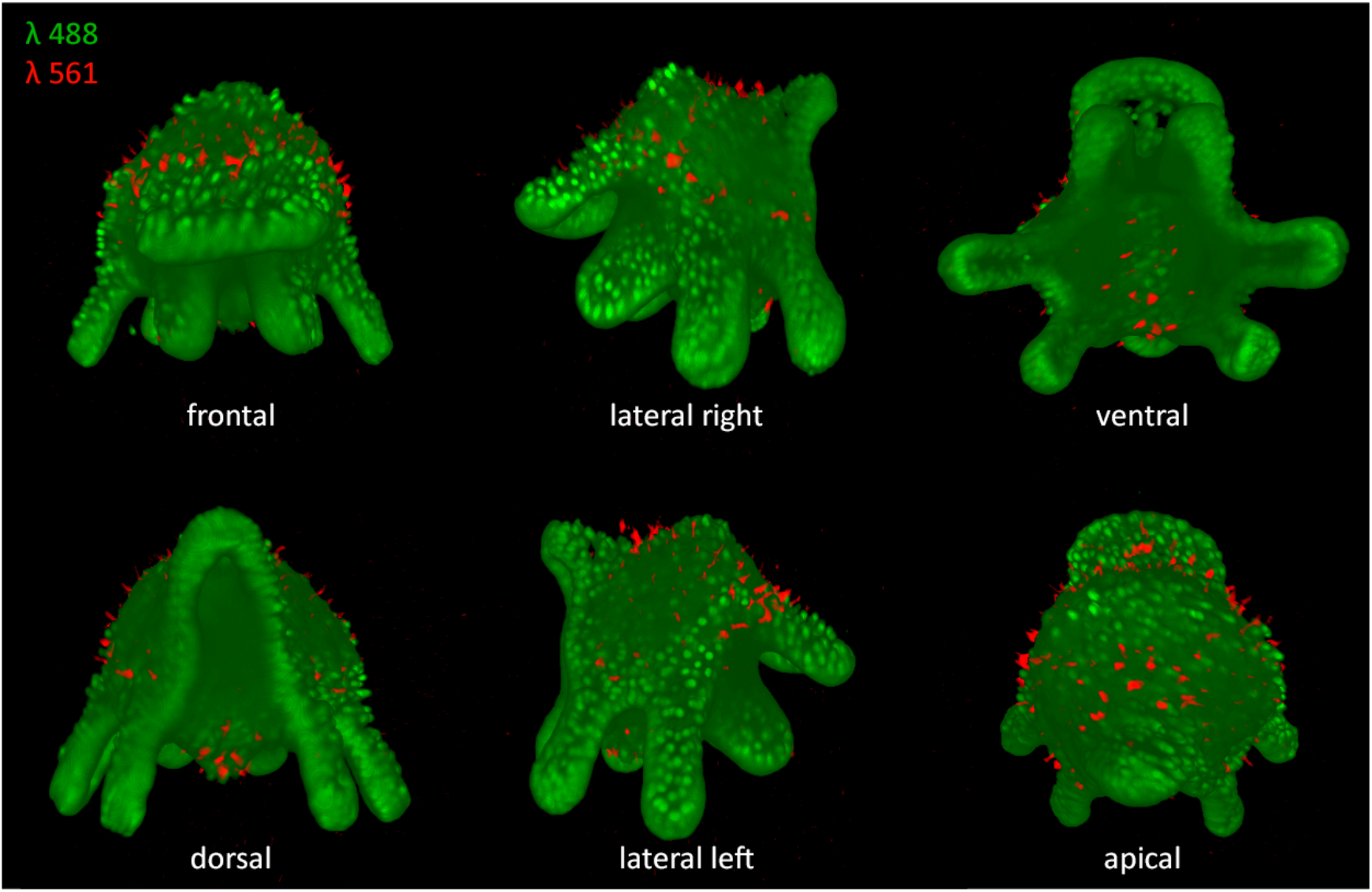
Several angles of a 3D-rendered Müller’s larva showing nuclei in green (captured with the 488 laser detection channel) and gland cells in red (captured with the 561 laser detection channel).

### OpenSPIM image acquisition with hardware controlled laser triggering is more than twice as fast

With the aim of reducing image acquisition time, we incorporated ESio’s TTL controller box (http://www.esimagingsolutions.com/) into our OpenSPIM microscope; this enables hardware-controlled synchronization of the timing of camera exposure and laser triggering. To test our ESio TTL controller, a 100 μm thick single color stack was imaged (1280x1080 resolution in 16-bit and an exposure time of 32 ms) with a constant step size of 1.5 μm. In this test, the software-controlled image acquisition (by MicroManager without the TTL controller box) completed the acquisition in 43.5 sec. In comparison, when hardware-controlled imaging is used, where lasers are triggered with the TTL controller box from ESio, image acquisition took 17.5 sec demonstrating a significant reduction of image acquisition time.

### Fiji’s bead based registration algorithm and multi-view deconvolution is essential to visualize all nuclei in *M. crozieri* embryos along the animal-vegetal axis

Having the possibility to 3D rotate the specimen and using a faster image acquisition set-up opens up for the possibility to carry out whole-embryo time-lapse videos of the development of an embryo. Fiji’s bead based registration algorithm and multi-view deconvolution plugins [4, 6] make it possible to fuse and deconvolve z-stacks imaged at multiple angles, acquired sequentially at any given time-point. We imaged the early development of *Maritigrella* covering the spiral cleavage and the formation of the four quadrants (Figure 8 A-J^TL^) and 3D-reconstructed a series of time-points using fixed embryos of several stages stained with Sytox-Green. To ensure that the development is normal in our live-imaging experiment, we created a series of cleavage stages with fixed specimens, for which both 3D reconstructions of immunostained OpenSPIM imaged embryos and SEM imaged embryos were performed (Figure 8 A-J^3D^ & A-J^SEM^). The 3D-reconstructed series of fixed embryos of several stages are available as 3D models (see Video 2).

### Rapid *in vivo* time-lapse sequences captured with OpenSPIM show the dynamic early embryonic development of *M. crozieri*

With the OpenSPIM equipped with two illumination paths and capable of rapidly producing, high-quality image stacks, we aimed to visualize the embryonic development in *M. crozieri* up to the 128-cell stage to further test the potential of OpenSPIM for live-imaging and, ultimately, for lineage tracing. We created a time-lapse sequence of a developing embryo injected at the one cell stage with a nuclear marker (H2B:GFP) and a membrane marker (CAAX:GFP). Our time lapse covers 18 hours and shows the stereotypical spiral cleavage and formation of four quartets and further development in *M. crozieri*. The sequence visualizes the embryo from the animal pole (Figure 8 A-J^TL^ and Video 1) and consists of 273 individual time-points captured approximately every 2 min. Development was not notably different between fixed and live specimens.

**Figure 8.**
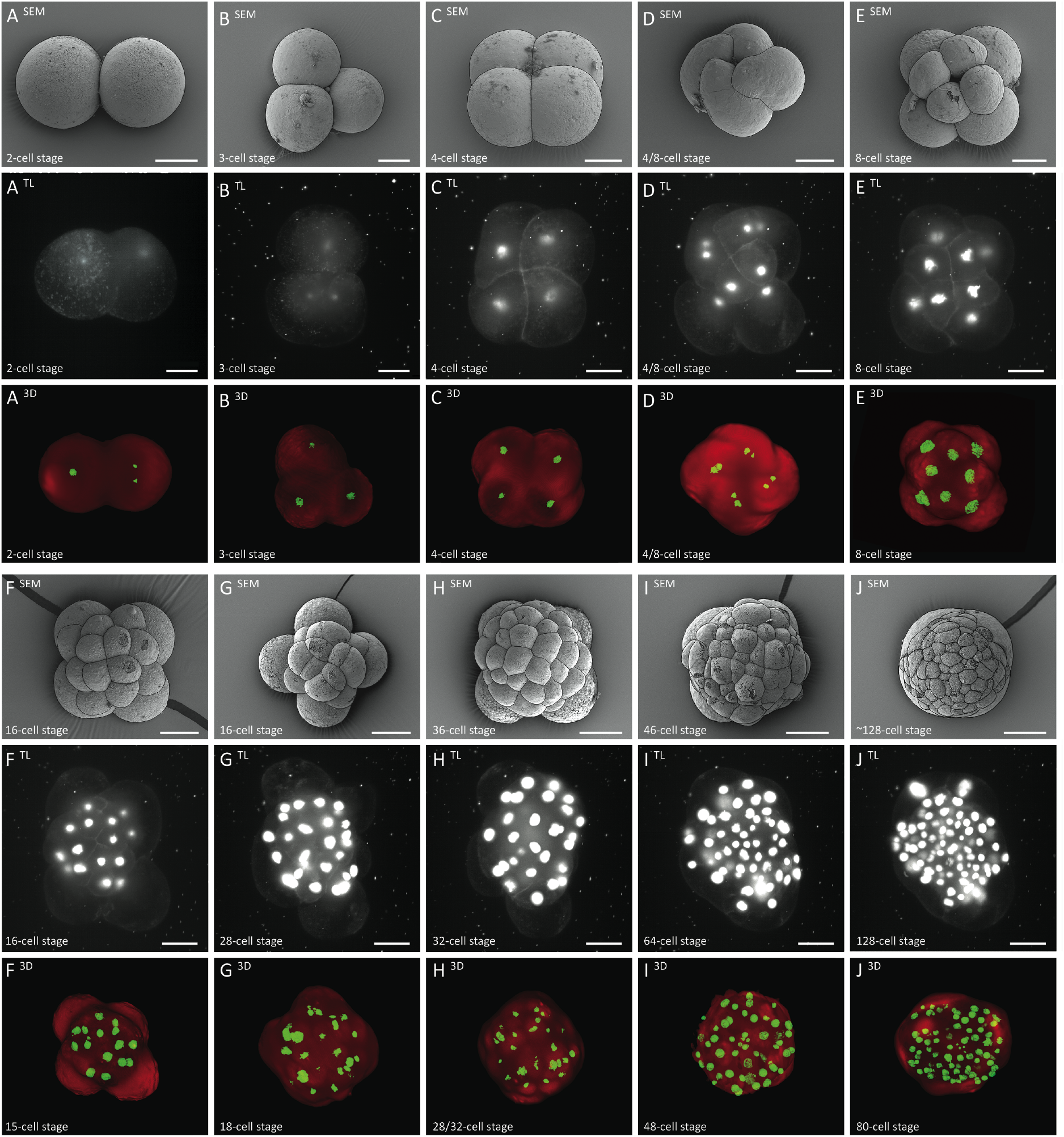
Summary figure of the early embryonic development of the polyclad flatworm *M. crozieri* (1-128-cell stages) (A-J); SEM pictures (A-J^SEM^) have been captured for comparison, stills from time-lapse sequences (A-J^TL^), multi-view 3D reconstructions (A-J^3D^); all embryos are shown from animal side. Note that time-lapse images (A-J^TL^) are presented as captured by the OpenSPIM (mirror images) and therefore cleavage direction is opposite to 3D and SEM images. All scale bars are 50 µm.

## DISCUSSION

### Summary of the capacity of our new OpenSPIM

#### New modifications tested for OpenSPIM

When designing microscopes for *in vivo* imaging with the purpose of tracing cells, one of the goals is to have a high imaging speed in order to have the best time resolution. One of our own modifications included **hardware-controlled imaging** that appears to be an elegant way of removing unwanted delays during image acquisition. Another relatively new implementation, at least in the context of OpenSPIM, is the use of **dual-sided illumination**, which we tested on our specimens. We demonstrated that bringing the light-sheet simultaneously from two sides to our opaque and yolky samples results in a significant increase in signal across the sample as shown for the nucleic markers of stained embryos (Figure 6).

### Points for consideration before purchasing and building an OpenSPIM

There is little doubt that a self-built light-sheet microscope is significantly more affordable than existing commercially available alternatives. The question that a laboratory considering whether to embark on building one ought to consider rather depends on two factors. First is the question of whether the finished microscope will be an adequate alternative in terms of image quality and ease of use for the specific task. Second, it is essential when considering an OpenSPIM to factor in the hidden costs involved, most obviously the costs implied by the time spent building the microscope, learning to use it and learning to use the open source software required to run the microscope and to process the data acquired.

For our purposes the quality and speed of acquisition that we were able to achieve with our home-built OpenSPIM device provides valuable, high quality data that suit our requirements. However, despite the fact that the assembly of an OpenSPIM is indeed quite straightforward, this step remains only one of the many challenges to overcome. Here we summarize various steps (Figure 9) we feel are worth considering before building an OpenSPIM in order to avoid assembling an expensive toy that will be forgotten shortly after.

**Figure 9.**
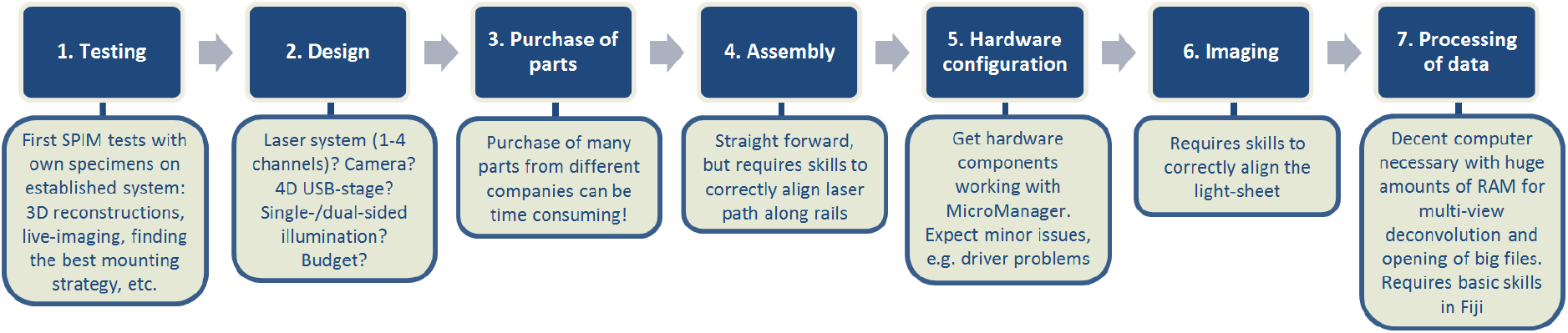
Flow chart illustrating steps necessary for establishing a home-built OpenSPIM

### Before you begin

Before beginning we would recommend prospective OpenSPIM users to image your own specimens on an established OpenSPIM system. This will provide valuable information on whether the system will be suitable for your purposes as well as show what is required in terms of hardware for capturing high-quality images of your particular specimens. This is also an opportunity to gain skills such as correctly aligning the light-sheets, getting familiar with the acquisition software, finding the optimal mounting strategy for the specimens and will inform decisions for the OpenSPIM design selected, as discussed in the next section. There are many OpenSPIM systems around the world; the current estimate is 70. The system from Tomancak lab also regularly travels to practical courses and was extensively used during the EMBO course on Light-sheet Microscopy in Dresden (2014 http://openspim.org/EMBO_practical_course_Light_sheet_microscopy and upcoming August 2016).

### Designing your OpenSPIM

The basic **design** of an OpenSPIM can be taken from the open access platform (http://openspim.org). However, modifications, which might meet more specific needs of individual users, require well thought-through decisions especially considering that most divergences from the basic plan will involve higher costs and, very likely, additional trouble shooting.

We have discussed the most significant amendments we have made in building our own OpenSPIM. Dual sided illumination allows us to image our relatively opaque, yolky embryos optimally and we have shown the benefits of this. Twin lasers allow us to observe more than one labelled molecule per sample. As the most expensive component, the laser system is of particular importance, especially when it comes to multi-channel acquisition. Our OpenSPIM can be easily upgraded up to as many as 4 different laser wavelengths whose beams are aligned within the laser system itself. This alignment of two lasers has proven robust even when travelling, meaning the microscope is fairly portable.

### Time taken for purchasing

While this might sound trivial, we found that purchasing elements occupied a significant time. There were three reasons for this, first that there is a multitude of suppliers to be negotiated with. Second, as some of the parts are expensive we (like many institutions) were required to obtain multiple quotes for each item. Third, the choices made regarding some parts had knock-on effects regarding the choice or specification of other parts. Finally it should not be forgotten that some of the parts of the OpenSPIM are bespoke and require a workshop or manufacturer for their production.

### Assembly

The **assembly** of all parts can be considered fairly easy, also thanks to the information provided from the http://openspim.org website. Usually this should be the least time consuming (and the most fun) task.

### Software and hardware integration

The MicroManager software and information provided on the OpenSPIM website makes correct **hardware configuration** relatively easy. However - at least in our experience - establishing the correct links between hardware components and the acquisition computer causes some time consuming problems. Additional time for hardware testing and configuration should be allowed. As an example, we experienced a major issue installing a simple FTDI chip driver, which is necessary for the ESio’s TTL controller box to communicate with the acquisition computer. Solving this problem required additional testing of the hardware and interaction with the original suppliers. Moreover, we strongly recommend interacting with the growing OpenSPIM online community via the mailing list, since many users are experiencing the same problems and it is the power of this community that will help you overcome them. Besides, hardware software integration is not something that a typical biologist can master with ease. Involving computer scientists or engineers on the undergraduate level, which should be relatively easy at any large University, is likely to smooth many integration problems. It will also give the students valuable experience with open access hardware and electronics and connect them with the active online communities in these areas.

### A fast way to correctly align the OpenSPIM

Learning how to correctly align the light-sheet is a skill that might require some help, but it is not particularly difficult to learn. Within the materials and methods we describe what we learned and how we currently align our two excitation light-sheets by simply adjusting the 25 mm and 50 mm telescope lenses and the two adjuster knobs (Horizontal & Vertical) of the Gimbal mounts of each corner mirror. It is a relatively fast approach (and certainly not the only one), but in our experience the achieved results in terms of image quality are more than satisfying.

### Image processing

Imaging and processing and the challenge of multi-view 4D microscopy of acquired data can be straightforward or become a major issue depending on the operator’s ambitions. Acquisition of z-stacks of fixed specimens and subsequent processing with Fiji can be easily learned and more sophisticated processing such as multi-view deconvolution can be learned using online Fiji tutorials on how to use the necessary plugins.

Considerably more challenging, in our experience, is long-term multi-view 4D microscopy of live embryos. This live-imaging setup (keeping the embryo alive and developing normally during acquisition for example) is clearly important. It is also essential to consider the challenge of post-processing the huge amount of data that are generated. Home-built OpenSPIMs are in principle capable of creating elaborate multi-view 4D microscopy vids on difficult specimens (e.g. opaque embryos with scattered emission-light). This was successfully demonstrated on *Drosophila* embryos, where data from six angles per time point have been acquired and the OpenSPIM data generated were subsequently successfully reconstructed using Fiji’s bead based registrations algorithm and fused via multi-view deconvolution. However, multi-view 4D microscopy requires an efficient flow of data-saving of the data produced onto hard drives. The second major requirement is a precise 4D motor system to keep the positional information of the specimen over time as exact as possible in concert with acquisition software that allows the correction of minor drifting of the specimen. Finally major computational resources are needed for processing the data generated, especially if multi-view deconvolution of hundreds of time-points is intended. In our opinion multi-view 4D microscopy is one of the most demanding and challenging experiments one can undertake with a home-built OpenSPIM and will thus be discussed further in the following section.

### Inefficient data saving can prolong time-point intervals during imaging

Multi-view 4D microscopy requires software that reliably saves large amounts of data on hard-drives without running into the problem of a data bottleneck. In single-view time-lapse videos, we observed that the creation of closely spaced time-points (intervals from about 90 sec / time-point) with the MicroManager SPIMacquisition plugin (available at OpenSPIM.org) can cause delays after a certain amount of time has passed. The interest here is, perhaps, less in the specifics of this issue and more in the observation that running an OpenSPIM (as opposed to a commercial system) will require the operator to get involved in many such technical challenges. Alternatively, as this is clearly a solvable issue, one could invest in collaboration with software engineers to identify the problem and adjust the open source software accordingly. Expert help from microManager, Fiji and OpenSPIM communities is expected and required. To ensure the problem is solved one has to invest in the solution. These communities are not compensated for developing the resources and their ability to fix specific problems is limited.

### Combining multi view acquisition with long term *in vivo* experiments

In our experience, long term *in vivo* multi view experiments with the aim of acquiring many time-points with several angles is not an easy task. Our living embryos occasionally undergo dynamic developmental processes, which can cause minor drifts during imaging. Additionally the automated correct positioning for each angle relies on the smooth and precise running of the USB-4D stage motors system (x, y, z, and twister motors) and on advanced acquisition software. Recently anti-drift plugins have been developed and implemented into MicroManager and are currently being further improved. We anticipate that these developments will bring major benefits for multi-view multi time point acquisition with an OpenSPIM.

### Processing of acquired multi-view data is challenging

The creation of multi-view 4D videos with an OpenSPIM has many interesting challenges. One important question that remains is how to deal with the huge amount of data generated. The processing of single time-points is feasible on a decent desktop computer (our system information can be found in the appendix), keeping in mind that SPIM registration processes such as multi-view deconvolution can require up to 128 GB of memory to successfully deconvolve a single time-point without compromising image quality and depending on parameters such as z-stack size, resolution, bit-rate, etc. To handle hundreds of time-points, even when imaging quality standards are lowered, a cluster computer with a sophisticated pipeline to organise the processing becomes a necessity. Cluster processing will certainly get more accessible in the future and an automated workflow for multiview SPIM recordings, see [17], but the need to set up such a pipeline and to have access to a cluster computer should also be borne in mind if OpenSPIM multi-view 4D microscopy is required.

## CONCLUSION

We have described the design and assembly of a T-configuration OpenSPIM with twin lasers. We have shown that a home-built SPIM microscope can be used as a scientific instrument to study the embryonic development of the polyclad flatworm *M. crozieri* on fixed specimens and *in vivo*. With our microscope we have produced high-quality 3D images of fixed larvae and have captured in detail the early embryonic development up to the 128 cell stage in a series of 3D reconstructed time-points.

One of our major goals is to use OpenSPIM for 4D microscopy (3D time lapse). Our OpenSPIM time-lapse videos, presented in Figure 7 and Supplementary Video 1, demonstrate our ability to image the embryogenesis of *M. crozieri* in unprecedented detail over time by single-view stack acquisition.

We have highlighted the problems encountered at all stages of building our OpenSPIM in the hope that this will help future users with similar ambitions. Building our microscope has been a fascinating challenge, and we conclude that OpenSPIM is eminently possible for anyone interested having continuous access to their own light-sheet microscope.

## Acknowledgements

We would like to thank Peter Brunt (from Laser 2000 (UK) Ltd) for his support with the multiple laser system (VersaLase) and the whole Tomancak lab at the MBI-CBG in Dresden and in particular Christopher Schmied for the valuable SPIM processing tips and help during our visits. We also want to thank Anna Czarkwiani for her input during writing the paper and Paola Oliveri for helping us setting up the microinjections at UCL.

## Funding

The OpenSPIM parts were principally purchased using the Biotechnology and Biological Sciences Research Council grant (BB/H006966/1). JG was funded by the Marie Curie ITN ‘NEPTUNE’ grant (no. 317172), under the FP7 of the European Commission. A.Z. was supported by the European Research Council (ERC-2012-AdG 322790) and F.L. by the Biotechnology and Biological Sciences Research Council grant (BB/H006966/1). P.T., M.H.-T. and P.P. were supported by The European Research Council Community’s Seventh Framework Program (FP7/2007- 2013), grant agreement 260746. M.J.T. is supported by a Royal Society Wolfson Research Merit Award.

## Competing interests

The authors declare that they have no competing interests.

## Author contributions

MJT and JG designed the experiments. PGP, PT, JG and FL designed the OpenSPIM. MHT helped JG establishing live-imaging and a microinjection setup for *M. crozieri* animals. JG and PGP built the OpenSPIM. JG and FS sampled and cultured experimental animals. JG obtained and injected embryos from gravid animals. JG performed OpenSPIM live-imaging and 3D reconstructions. AZ and JG made SEM images. JG prepared the figures. JG and MJT analyzed the data and wrote the manuscript draft. PT and MHT contributed to the writing. All authors read and approved the final manuscript.

## Movie Legends

**Video 1** – Left: 18 hours of continuous live imaging of the embryogenesis of M. crozieri by single-view stack acquisition (live staining achieved by mRNA injections: CAAX-GFP marking the membranes and H2B-GFP marking the nuclei). Cell stages (defined by manually counting the nuclei) are shown at the top left corner in red. Time at the top right corner (cyan) indicate hours post oviposition (hpo). Scale bar = 50 um. Right: Selected scanning microscopy pictures, which correspond to live-imaging stages.

**Video 2** – (A-J) 3D-models of a series of fixed embryos of several stages reconstructed using Fiji’s bead based registrations algorithm and multi-view deconvolution.

## Supplementary Figures

**Supplementary Figure 1.**
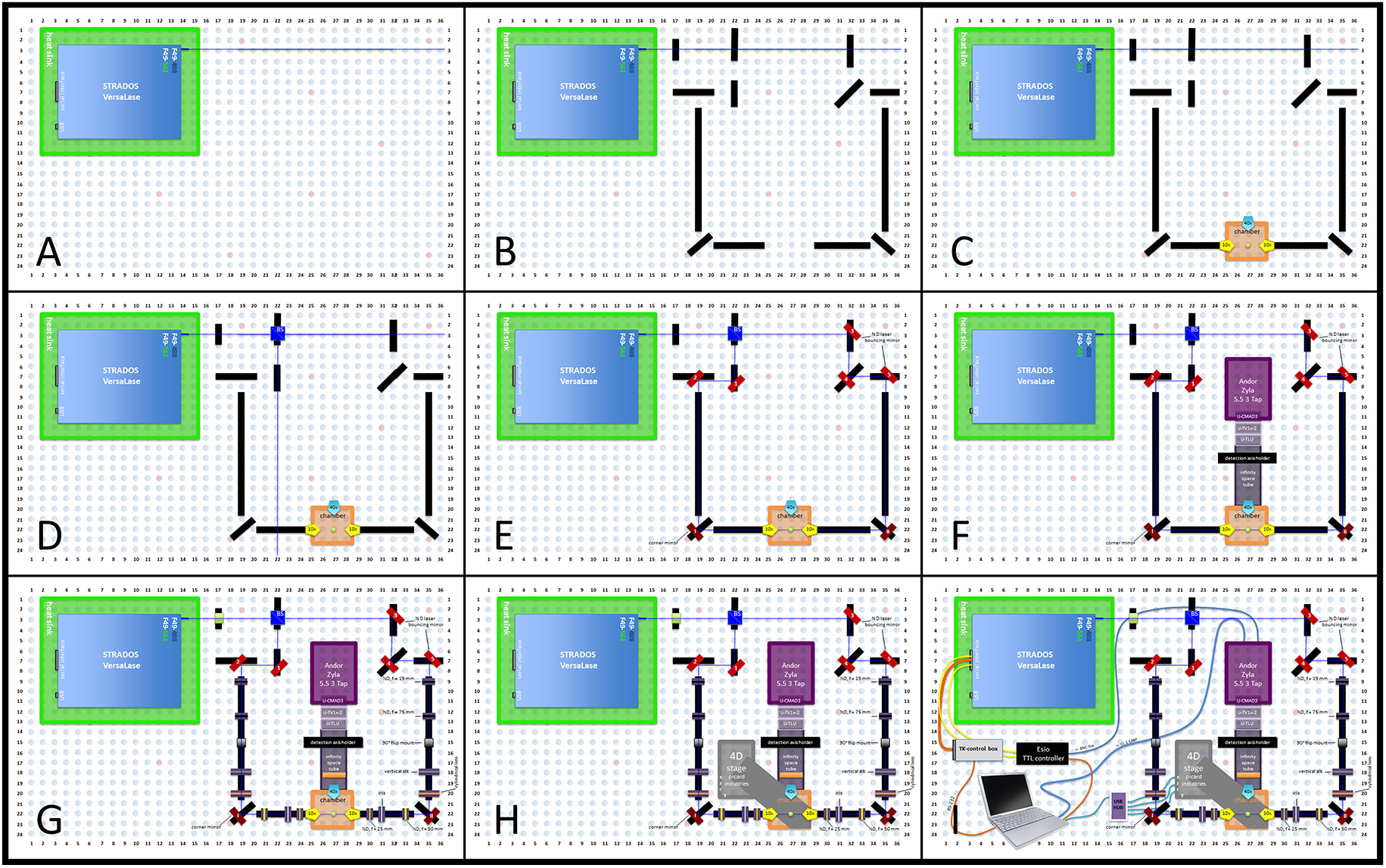
Schematic assembly of the OpenSPIM; **(A)** Step 1 - Installation of breadboard feet onto the optical breadboard and placing it on its final position; Step 2 - Installation of laser heatsink on the optical breadboard and fixation of laser system (VersaLase) on top of the heatsink **(B)** Step 3 - Cutting of optical rails for corner mirrors and two reflecting mirrors and installation of rail system onto the optical breadboard **(C)** Step 4 - Installation of the pre-assembled acquisition chamber onto the corresponding rail **(D)** Step 5 - Installation of the beam splitter **(E)** Step 6 - Installation of all corner and laser reflecting mirrors **(F)** Step 7 - Installation of detection axis holder, infinity space tube, camera and its corresponding connection adapter units to the infinity space tube (U-CMAD3, U-TV1x-2 and U-TLU) **(G)** Step 8 - Installation of optical elements (beam expanders, telescope); Step 9 - Installation of clean-up and emission filters **(H)** Step 10 -Installation of Picard 4D stage on its correct position **(I)** Step 11 - Plugging in the controller boxes (Esio TTL controller box & VersaLase control box), VersaLase, Camera, USB 4D-stage and connecting them up with the acquisition computer

**Supplementary Figure 2.**
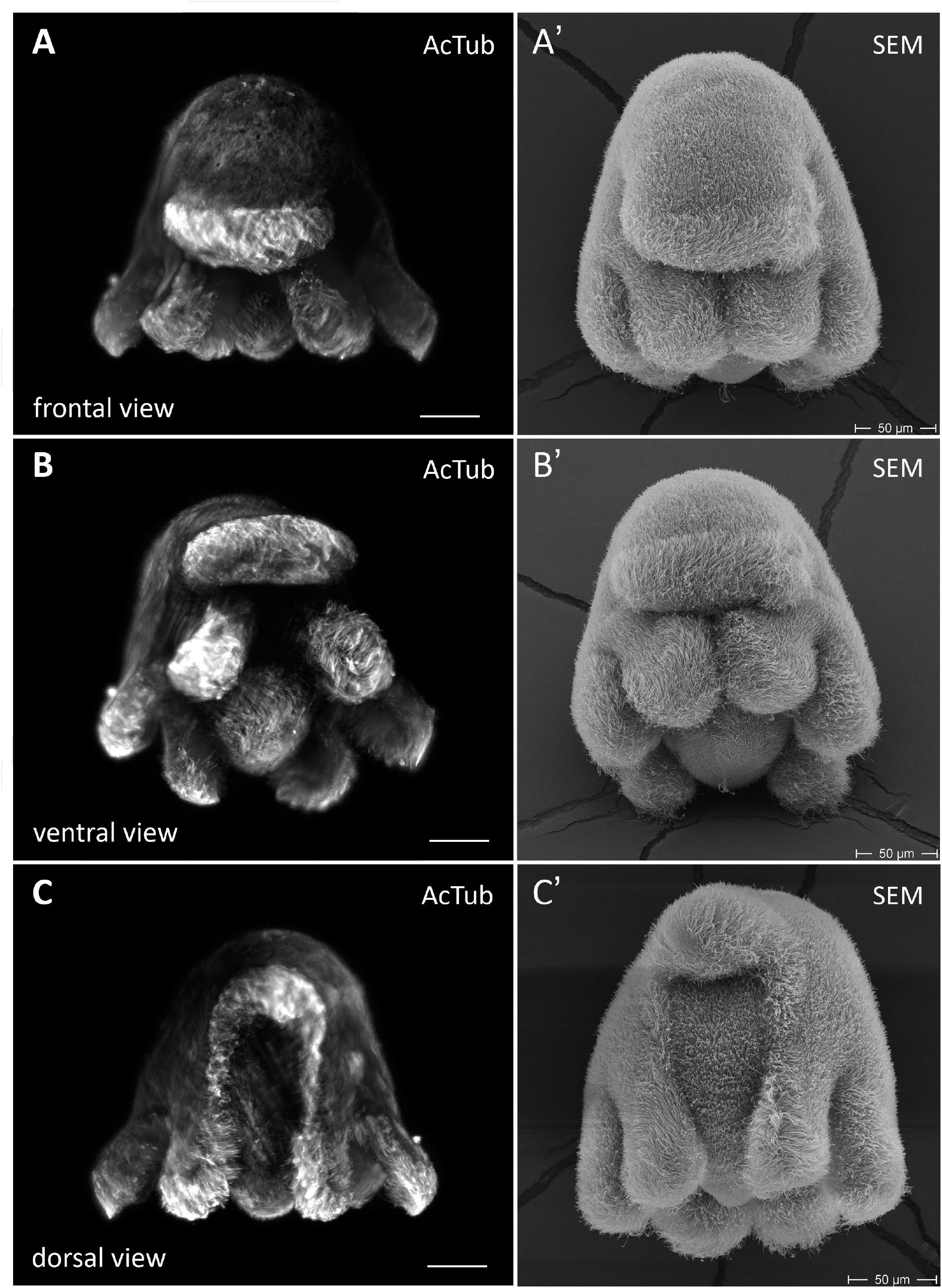
Images (maximum projections) of fixed Müller’s larvae stained with Acetylated tubulin and captured with our OpenSPIM images show a clear resemblance to scanning electron microscopy images of similar stage larvae

